# Threshold Selection for Brain Connectomes

**DOI:** 10.1101/2021.10.09.463759

**Authors:** Nicholas Theis, Jonathan Rubin, Joshua Cape, Satish Iyengar, Konasale M. Prasad

**Affiliations:** Department of Psychiatry, University of Pittsburgh School of Medicine, Pittsburgh PA; Department of Mathematics, University of Pittsburgh, Pittsburgh PA; Department of Statistics, University of Wisconsin–Madison, Madison, WI; Department of Statistics, University of Pittsburgh, Pittsburgh PA; VA Pittsburgh Healthcare System, Pittsburgh, PA; Department of Bioengineering, University of Pittsburgh Swanson School of Engineering, Pittsburgh PA

**Keywords:** Networks, Graph theory, Magnetic resonance imaging, Biomedical imaging, Network Thresholding

## Abstract

**Introduction:** Structural and functional brain connectomes built using macroscale data collected through magnetic resonance imaging (MRI) may contain noise that contributes to false-positive edges, which can obscure structure-function relationships with implications for data interpretation. Thresholding procedures are routinely applied in practice to optimize network density by removing low-signal edges, but there is limited consensus regarding the appropriate selection of thresholds. We compare existing methods and propose a novel alternative objective function thresholding (OFT) method.

**Methods:** The performance of thresholding approaches, including a percolation-based approach and an objective function-based approach, is assessed by (a) computing the normalized mutual information (NMI) of community structure between a known network and a simulated, perturbed networks to which various forms of thresholding have been applied, and (b) comparing the density and the clustering coefficient (CC) between the baseline and thresholded networks.

**Results:** In our analysis, the proposed objective function-based threshold exhibits the best performance in terms of high similarity between the underlying networks and their perturbed, thresholded counterparts, as quantified by NMI and CC analysis on the simulated functional networks.

**Discussion:** Existing network thresholding methods yield widely different results when graph metrics are computed. Thresholding based on the objective function appears to maintain a set of edges such that the resulting network shares the community structure and clustering features present in the original network. This outcome provides proof-of-principle evidence that thresholding based on the objective function could offer a useful approach to reducing the network density of functional connectivity data.

**Impact Statement:** Network thresholding refers to removing edges between node pairs in a functional network that have weak edge-weights that may arise from unwanted variability or noise. Since edge-weight cutoffs used to generate a binary network can be sensitive to thresholding, we introduce a novel thresholding algorithm. We find that when applied to networks derived via perturbations, namely through simulated functional connectivity of a known network, this approach yields a binary network that is more similar to the known network compared to using existing thresholding approaches. Thus, our algorithm is a competitive candidate for use in thresholding of brain connectome.

## INTRODUCTION

Network representations of biological data often produce graphs that are weighted but fully connected, consist of low-signal edges (van den Heuvel et al., 2017), or are sparse but contaminated with false positives (Drakesmith et al., 2015, Pu et al., 2015). For instance, a functional connectome (FC) generated from functional MRI (fMRI) data consists of edges that are defined by correlations between nodal time series of blood oxygenation level dependent (BOLD) signals. Weighted networks can include weakly correlated regions that may represent noise. In the structural connectome (SC) built using anisotropy-based diffusion streamline, false positives are less prevalent but still possible (Sotiropoulos and Zalesky, 2019).

Network thresholding is crucial to elucidate differences between study groups in a reproducible manner. A threshold can be applied to create less dense binarized or weighted graphs from fully connected graphs (Bassett et al., 2008), or to remove false positive observed edges (Drakesmith et al., 2015). Nevertheless, thresholding procedures have limitations. Aggressive thresholding approaches produce graphs that are too sparse, whereas overly cautious approaches produce graphs containing noise-driven connections, impugning the accuracy of subsequently calculated graph metrics. For real-world applications of graph theory such as in psychiatry, these issues confound the interpretation of differences in graph measures between clinical groups. Despite the importance of thresholding, few comparisons among existing methods have been made in the literature to evaluate thresholding techniques.

There are several difficulties in comparing thresholding techniques. Existing thresholding methods sometimes apply in distinct settings or give different types of information. Some earlier thresholding methods, such as the small-worldness (σ) range (Bassett et al., 2008), provide a range of thresholds, but not a specific optimal threshold. The σ-range approach also requires that the networks themselves have small-world properties, thereby limiting its application. Statistical thresholding (Chen et al., 2008, Ferrarini et al., 2009) requires that edge weights have associated metadata, such as *p*-values for correlations, that are not attributed to all biological networks, and ultimately also require the researcher to select a cutoff value for statistical significance. For example, SC networks constructed using diffusion stream counts do not have an associated *p*-value. More recent methods rely on downstream classifier accuracy to detect the best network threshold (Drakesmith et al., 2015, Zanin et al., 2012), e.g., the “multi-threshold permutation correction” method (Drakesmith et al., 2015) involves determination of an optimal threshold by clustering thresholded networks into groups and evaluating which threshold yields the most accurate classification. While these methods are applicable to networks with a variety of structures (i.e., not just small world), they cannot be performed on a single graph, or a collection of graphs from a single or multiple unknown groups.

The percolation threshold, which is defined as the highest threshold value for which a network’s giant connected component (GCC) includes all nodes, circumvents these difficulties. This is a straightforward and reasonable selection method, but it does not apply to networks that are not fully connected without any thresholding, i.e., the GCC of the network is not the full set of nodes (Bordier et al., 2017). For example, in resting state fMRI many nodes might not show significant activation and in task-fMRI, some nodes may not participate in processing a task.

In addition, it is difficult to define a universal performance measure for outcomes of thresholding. The normalized mutual information (NMI) is useful to compare different thresholding methods. The NMI provides a comparison between the community structures identified within any pair of networks defined on the same set of nodes, each featuring at least two communities (Alexander-Bloch et al., 2012). If both networks being compared have been subject to thresholding, then this comparison is problematic (e.g., making both networks very sparse might yield a high NMI, but based on communities that bear little resemblance to those present in the true positive edges in the original graphs). Such a comparison is also not possible if the graphs do not feature community structure.

To overcome these challenges, and to offer a more universally applicable, easy to implement option for thresholding, we introduce a method based on the calculation of an objective function that uses a simple optimization approach to compute an edge weight threshold for *any* given network, calculated based on any measure of a property of a graph related to the edges that it includes, or graph metric, such as characteristic path length (λ). The objective function is based on computing a graph metric over a range of reasonable thresholds, chosen to yield thresholded graphs that are connected but have graph density smaller than one. The selected threshold yields an extremum of an objective function that we define, representing maximal deviation of the graph metric from its value at the problematic extremes of the threshold range, which are where the graph’s density is too high, or its GCC is too small. To demonstrate the performance of this method, we start from a computationally generated, ground truth network designed to exhibit a certain community structure (Lancichinetti A et al., 2008) and we introduce fMRI-like noise (Welvaert et al., 2011) to perturb the starting networks into fully connected networks inspired by functional networks. We compare the performance of the OFT with other thresholding methods when applied to these perturbed networks, in terms of their success in restoring the community structure, density, and clustering properties of the ground truth network.

## METHODS

### LFR Simulations

To evaluate the effectiveness of thresholding techniques, we generated a sample of networks using a published approach (Bordier et al., 2017). First, a benchmark Lancichinetti-Fortunato-Radicchi (LFR) (Lancichinetti A et al., 2008) network is produced, where the true community structure is known. LFR networks were generated using 363 nodes, which was selected because the Human Connectome Project (HCP) multimodal atlas has 360 nodes plus brainstem and left and right subcortex (Glasser et al., 2016). The LFR networks also require community mixing parameters to be specified, which were selected based on the guidance of previous use of the LFR benchmark algorithm for network neuroscience applications (Bordier et al., 2017). Next, fMRI-like noise (Welvaert et al., 2011) was introduced to each LFR network to produce a corresponding fully-dense (density=1) functional connectivity simulation (FC_sim_) network, with parameters based on previous work (Bordier et al., 2017). The following parameters were used for the LFR network based on prior work cited above: 363 nodes with an average degree of 24, maximum target degree of 107, community size minimum of 12 and maximum of 32 nodes, with both the community mixing and weight mixing coefficients set to 0.2. For the FC_sim_ parameters, a signal-to-noise ratio (SNR) of 35 was used and 100 replicates were examined to simulate the acquisition of 100 fMRI connectomes. Each replicate was generated from the same LFR network, so all FC_sim_ replicates share the same ground truth modular structure.

The effectiveness with which thresholding of the FC_sim_ replicates for noise removal recovers the properties of the benchmark LFR network could then be evaluated, across a variety of thresholding approaches. To do this, the NMI (see below) between the known community assignments of the original LFR network and the community structure detected in the noisy LFR networks after thresholding were compared among thresholding techniques (Bordier et al., 2017). Other measures, such as nodal density and clustering coefficients (CCs) of the binary networks, were also compared to the original LFR network for additional network-specific performance measures. **Figure 1** shows an overview of the experimental design.

**Figure 1:**
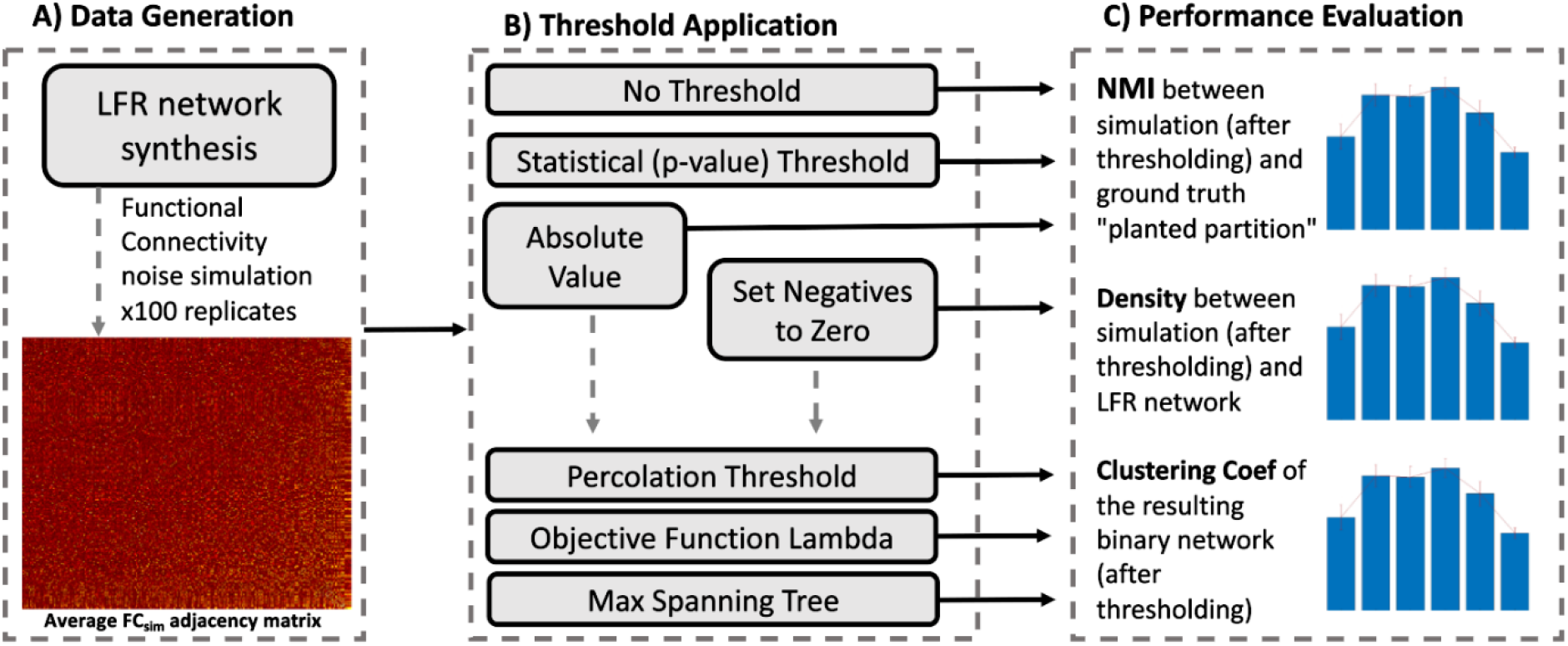
The experimental design for the study. A. Data generation consists of generating LFR network, then adding 100 replicates of functional MRI-like noise. B. Threshold application is preceded by either absolute value of the weighted FC simulated networks to preserve strong negative values, or by “zeroing out” the negative values under the assumption that negative edges overwhelmingly represent node pairs that are not directly connected in the ground truth LFR network. These considerations only apply to the percolation threshold, objective function, and maximum spanning tree. A fully weighted “no threshold” network is included as a “negative control”. C. Performance of the thresholding methods is compared to the original LFR network using NMI, density, and clustering coefficient.

### Threshold Space

Network thresholding provides a cutoff to convert a weighted network into a binary network consisting of presumed true positive edges, such that the original edges with weights at a given threshold, θ, are set to zero, while all other original edge weights are set to the value one. We use “threshold space” to refer to the set of threshold values tested when evaluating a range of thresholding choices; for example, a natural approach might be to consider a set of possible thresholds between two values θ_0_ and θ_1_, separated by a fixed step size ε, in which case our threshold space would be the vector of values (θ_0_, θ_0_+ε, θ_0_+2ε,…, θ_1_-ε, θ_1_), assuming that (θ_1_-θ_0_)/ε is a positive integer. A threshold space is not unique to a specific thresholding method, but it is a critical ingredient in a given method’s implementation. A “complete” threshold space would include every unique edge weight represented in a network, to the highest numerical precision possible. Since this choice is impractical in terms of computation time for large networks or large numbers of networks, we define a “rounded rank” threshold space. This is the set of unique edge weights, ranked lowest to highest, after rounding all weights to the nearest thousandth place.

### Ranked versus Combinatorial Thresholding

A distinction can be made between a “ranked” threshold space and a “combinatorial” thresholding method. Unlike a ranked approach, a combinatorial method can remove edges that have larger weights than some of the edges it preserves, e.g., statistical thresholding and maximum spanning tree (see below). Most thresholding methods operate on a ranked basis.

### Negative Edge Weights

In a network with some edges having negative weights, a positive threshold will force the zeroing out of all negative weights. In a network in which edge weights correspond to a count or level of a physical object, such as number of fiber tracks in structural networks, negative edge weights could result from noise or error and hence their removal is appropriate. In other cases, negative edge weights can pose a challenge for network thresholding, e.g., in a network with edge weights based on correlations; large negative values actually represent stronger relationships than near-zero negative values. In this situation, applying the absolute value function to all network edges before defining a threshold space could be an appropriate step to ensure that thresholding will preserve the strongly negative edges and not the weakly negative edges. We apply both approaches in our analysis.

### Graph Metric Calculations

Graph metrics, including density, CC, *λ*, modular structure, and the size of the GCC were calculated using the Brain Connectivity Toolbox in MATLAB (Rubinov and Sporns, 2010). For modular structure, the community detection algorithm implemented in the BCT relies on the Newman method (Newman, 2006) which is a spectral partitioning method that expresses this maximization problem in terms of a matrix eigenvector. The communities are determined by searching for the community structure that maximizes the modularity coefficient (Q).

In networks that have GCCs with fewer than 100% of the nodes, i.e., where disjointed sub-graphs appear, the calculation of *λ* treats the disconnected components as separate networks and averages their results, ignoring pairs of nodes that are not in the same component.

### Percolation Threshold

When a ranked threshold space is used, a larger threshold results in the removal of more edges. The percolation threshold is defined as the largest ranked threshold at which the network’s GCC contains all nodes in the network (Bordier et al., 2017), such that the network is connected. This threshold aims to find the optimal balance between information gained by noise removal and information lost by excessive pruning of edges, assuming that all nodes belong to the same network.

We will use the notation GCC=1 to indicate that 100% of the nodes in a network are in its GCC and the notation GCC=*x* for other *x* values between 0 and 1 to indicate the fraction of nodes in a network in its GCC when the network is no longer connected. The percolation method and the maximum spanning tree (MST) (see below) both provide binary networks with GCC=1; the former is the strictest possible “ranked” threshold to achieve this, and the latter the strictest possible “combinatorial” threshold.

### Objective Function Threshold

We developed the OFT method under the assumption that the optimal threshold for any weighted network lies between the smallest threshold for which the network density is not 1, and the largest threshold for which the GCC includes the experimentally determined percentage of nodes, here 100%. The objective threshold is determined by first employing a threshold sweep where a range of possible thresholds are evaluated. For a given graph measure, M, such as density, *λ*, transitivity, or any other scalar graph summary metric, the optimal threshold should maximize the difference of this metric from the value of the same metric at both end points of the threshold space, termed M_0_ and M_1_. The objective function can be extremized across this range to find the optimal threshold for a given graph metric in any weighted network (**Figures 2**). We tested a variety of graph metrics, including density, average nodal degree, *λ*, transitivity, average CC, and efficiency; *λ* was found to be the best option using the NMI criteria to evaluate performance (**Figure 3**). The nearest alternative, transitivity, was significantly lower from λ by a paired t-test (t=8.19, p<0.0001).

**Figure 2:**
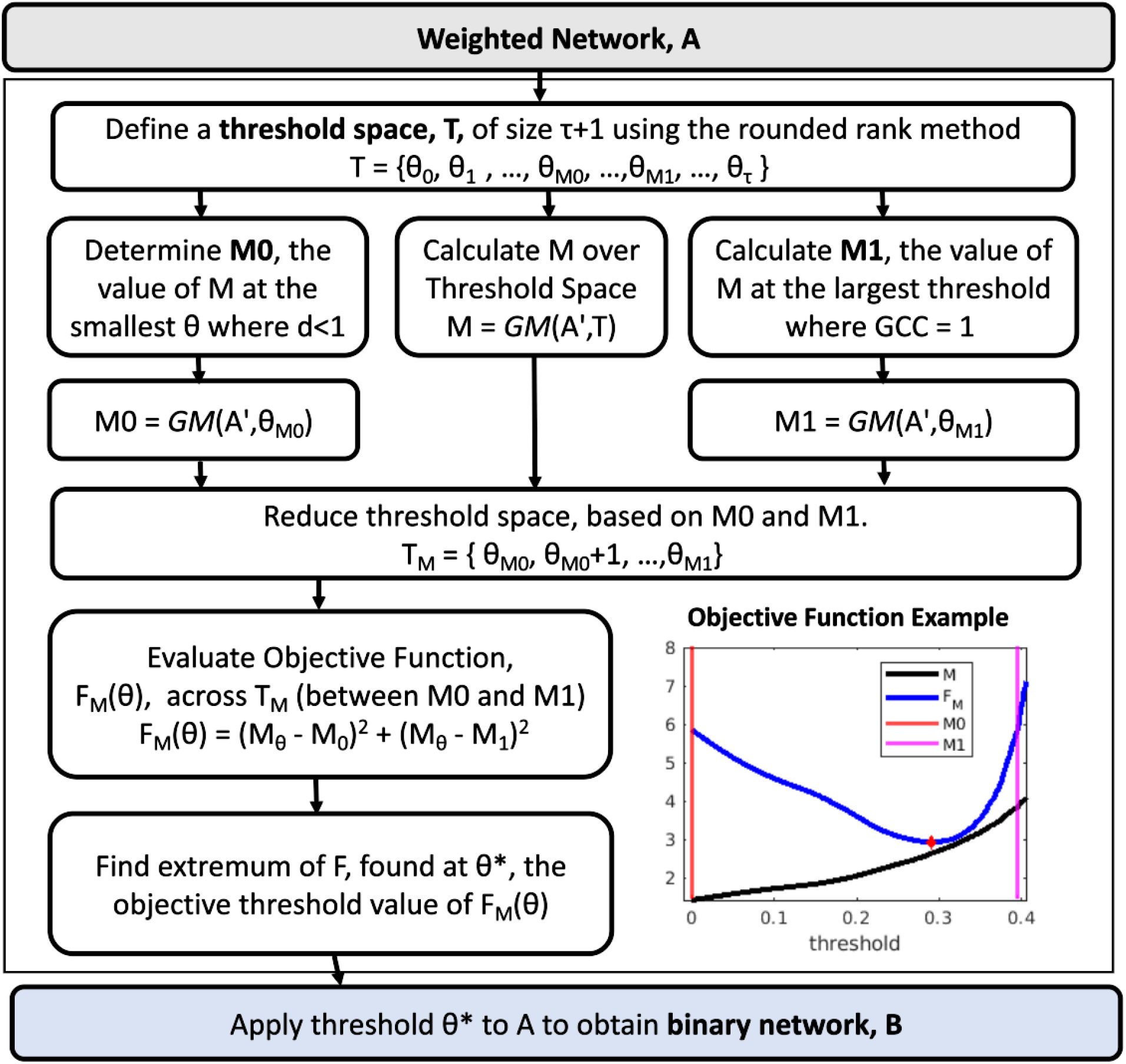
Overview of the Objective Function. Key terms: A, the weighted adjacency matrix; A’, the weighted adjacency matrix with negatives zeroed out, or after absolute value is taken; T, the threshold space; τ, size of threshold space; ϑ, any arbitrary threshold space; ϑ, any arbitrary threshold value; M, any graph metric (throughout this paper we use M=lambda); GM, the function that defines M, applied to A’ after binarization at threshold at ϑ; GCC, the giant connected component. If the GCC is less than 1, not all nodes are reachable. T_M_ represents a reduced threshold space, constrained by M0 and M1 conditions. The Objective Function, F, is defined for any given M and threshold space, and has an extremum at ϑ*, the objective threshold, which produces a binary network, B. Inset graph: a single network-level example of the objective threshold method evaluated for M = characteristic path length (λ).

**Figure 3:**
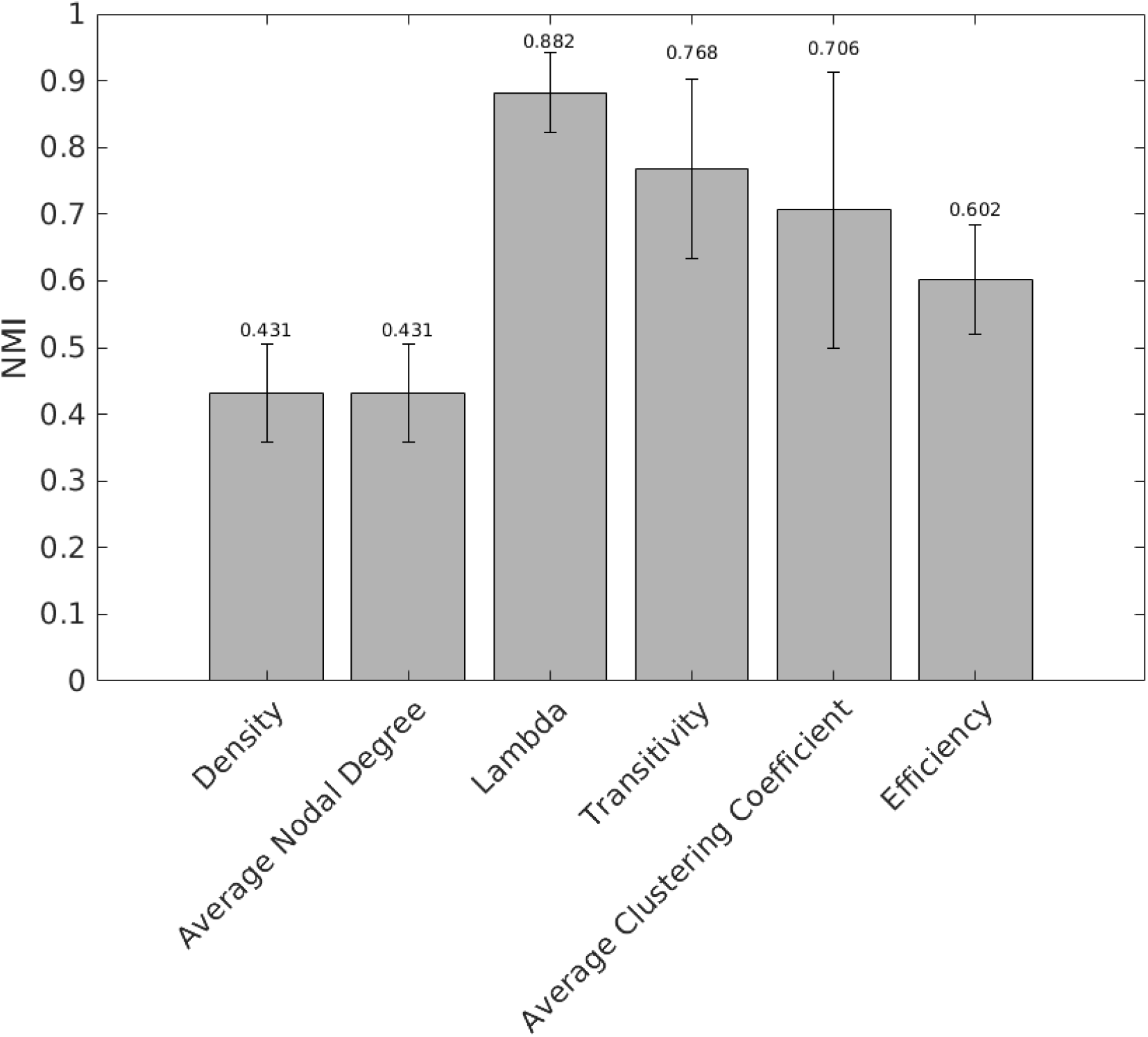
NMI was calculated between the LFR ground-truth community structure and the N=100 FC_sim_ networks after thresholding, using the objective function method with GCC=1 and rounded rank (to the nearest thousandth) threshold space, and one of six choice of graph measure (M). Absolute value of weighted networks was used, and removing negatives was not assessed for this comparison. Density, average nodal degree, λ, transitivity, average clustering coefficient, and efficiency were compared to identify the best choice of M in the objective function for this application. The highest NMI was achieved with lambda, and the nearest alternative, transitivity, was significantly lower from λ by a paired t-test (p<0.0001, t = 8.19).

The objective function takes the form:

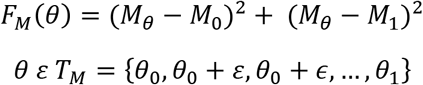

where *θ* is any threshold at which the function is being evaluated and *M* refers to the graph metric being used, with *M_θ_* denoting the value of *M* evaluated on a graph that has been binarized using threshold *θ* and *M_0_, M_1_* denoting the values of *M* at the extreme thresholds. Each threshold is taken from the threshold space *T_M_* of thresholds tested, ranging in steps of ε=0.01 from *θ_0_=kε* to *θi=Kε*, for integers *k<K*, such that the boundary thresholds *θ_0_*, *θ_1_* are both in the interval [0,1]. As noted above, *θ_0_* is selected as the lowest threshold that gives density <1, which for a sparse network is *θ_0_=0*. Moreover, we used the percolation threshold as the upper bound for our objective function calculation, although it is possible that larger choices of this bound (for which GCC<1) could also work well. That is, for a connected graph, such as the graph obtained by thresholding with the percolation threshold, all nodes are in the GCC. From there, if edges are removed uniformly at random, there is typically a rather abrupt transition from a GCC that includes all nodes to a much smaller GCC (Newman, 2001). Hence, very different GCC sizes will result from small increases in the threshold above the percolation threshold, and thus we would end up with similar *θ_1_* values by selecting the smallest threshold such that the proportion of nodes in the GCC is ≥x% for any choice of x between 100 and some much smaller values (approximately 20 in our empirical explorations). For biological networks that are known to comprise multiple components, a larger choice of *θ_1_* than the percolation threshold would clearly be appropriate, but such networks are not considered here.

### Optimization of OFT

We define the optimal threshold based on a given graph metric, *M*, from within the threshold space *T_M_*, based on the values of the objective function *F_M_*(*θ*) evaluated for all *θϵT_M_*. Specifically, we select the optimal threshold as the value at which *F_M_*(*θ*) has an extremum on the interval [*θ_0_*, *θ_1_*]. This criterion selects the threshold at which the value of *F_M_* is farthest from its endpoint values. Heuristically, this selection is based on the idea that at a sufficiently low threshold, weak edges are prevalent in the graph, to an extent that will likely have a strong influence on graph metric values. At a sufficiently high threshold, the network will have split (GCC<1) or be on the verge of splitting (GCC=1) into disconnected components, which likely corresponds to meaningful edges having been discarded, also likely to have a strong influence on graph metrics. Thus, a useful threshold would be one for which the graph metric of interest deviates significantly from its values at these endpoints.

In practice, we find that either *M_θ_* rises from a low value at or near 0 at *θ_0_*, has an interior maximum, and then declines back to near 0 again at *θ_1_*, or else *M_θ_* varies monotonically as *θ* increases from *θ_0_* to *θ_1_*. In the former scenario, the optimal threshold is simply the threshold in (*θ_0_, θ_1_*) where the maximum of *F_M_(θ)* occurs. In the latter case, we have *F_M_(θ_0_)=F_M_(θ_1_)=(M_0_-M_1_)^2^*. At the same time, if *F_M_(θ)* is monotonically increasing from *θ_0_* to *θ_1_*, then for any *θϵ(θ_0_, θ_1_), M_e_ < M_θ_ < M_1_*, and by the triangle inequality,

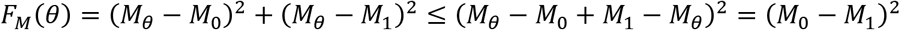

That is, *Fm_M_(θ)* takes its maximal values at *θ_0_* and *θ_1_* and has an interior minimum between these extremes, at the *θ* value where *M_θ_ = (M_0_ + M_1_)/2*. A similar argument applies and gives an interior local minimum of *F_M_(θ)* at the analogous *θ* value if *M_θ_* is monotonically decreasing from *θ_0_* to *θ_1_*.

### Statistical Threshold

Statistical thresholding involves retaining edges based on statistical significance and excluding edges that are not statistically significant with respect to a given measure (Bullmore and Bassett, 2011). We used a nominal uncorrected cutoff p<0.05 to determine whether a correlational edge should be included in the binary network or not. The statistical threshold can be considered as a combinatorial thresholding technique because it can remove edges that have a higher weight than some other edges it does not prune.

### Maximum Spanning Tree (MST)

Kruskal’s Algorithm (Kruskal, 1956), as implemented in a MATLAB package (MATLAB Central (Li, 2022), was used to determine the MST, which is defined as the smallest subset of edges in a graph that connects all nodes (GCC=1); specifically, this algorithm selects the spanning tree with the heaviest weighted edges possible, whence the “maximum” spanning tree nomenclature. Notably, while the MST for a graph has a fully connected component (GCC=1), unlike the percolation threshold that also yields a network for which the GCC=1, the MST does not include any redundant edges. Therefore, the MST thresholding method can be considered a combinatorial threshold because edges may be removed that are higher-weighted than other lower-weighted edges that are not removed.

### Normalized Mutual Information (NMI)

The NMI was calculated in previous publications as an evaluation criterion (Alexander-Bloch et al., 2012). Briefly, the NMI is a value between 0 and 1 that rates the similarity of two arbitrary lists of the same length, with 0 indicating completely chance correspondence, and 1 indicating perfect correspondence. NMI values near 0 represent limited shared information, whereas values near 1 represent substantial shared information. We compared two groupings of nodes into modules: that of the “ground truth” modular community structure for the LFR network, and that of each FC_sim_ network after thresholding. NMI thus quantifies the extent to which the module assignments for the nodes in each thresholded FC_sim_ network agree with the nodes’ assignments in the LFR network.

## RESULTS

We generated LFR networks to represent baseline networks with nonnegative edge weights, featuring an element of modularity, reminiscent of human brain structural connectome networks. From this ground truth LFR benchmark network, we applied fMRI-like noise, independently across 100 trials, resulting in FC_sim_ networks. For our purposes, the important features of these FC_sim_ networks are that they are perturbations of the original LFR networks, that all edge weights are non-zero, and that there are edges with negative weights. In the simulation context, we found that out of 65,703 possible LFR edges total in the off-diagonal, 4,082 are non-zero in the LFR network, i.e., dyadically connected in the LFR. Here 661 of these 4,082 (16%) are negatively weighted in the typical FC_sim_. On average, 26,740 FC_sim_ weights were negative, indicating that a randomly selected negatively weighted FC_sim_ edge had a 2.5% chance (661/26,703) of representing a true positive. The relationship between edge weights in the LFR and the FC_sim_ is depicted in Figure 4.

**Figure 4:**
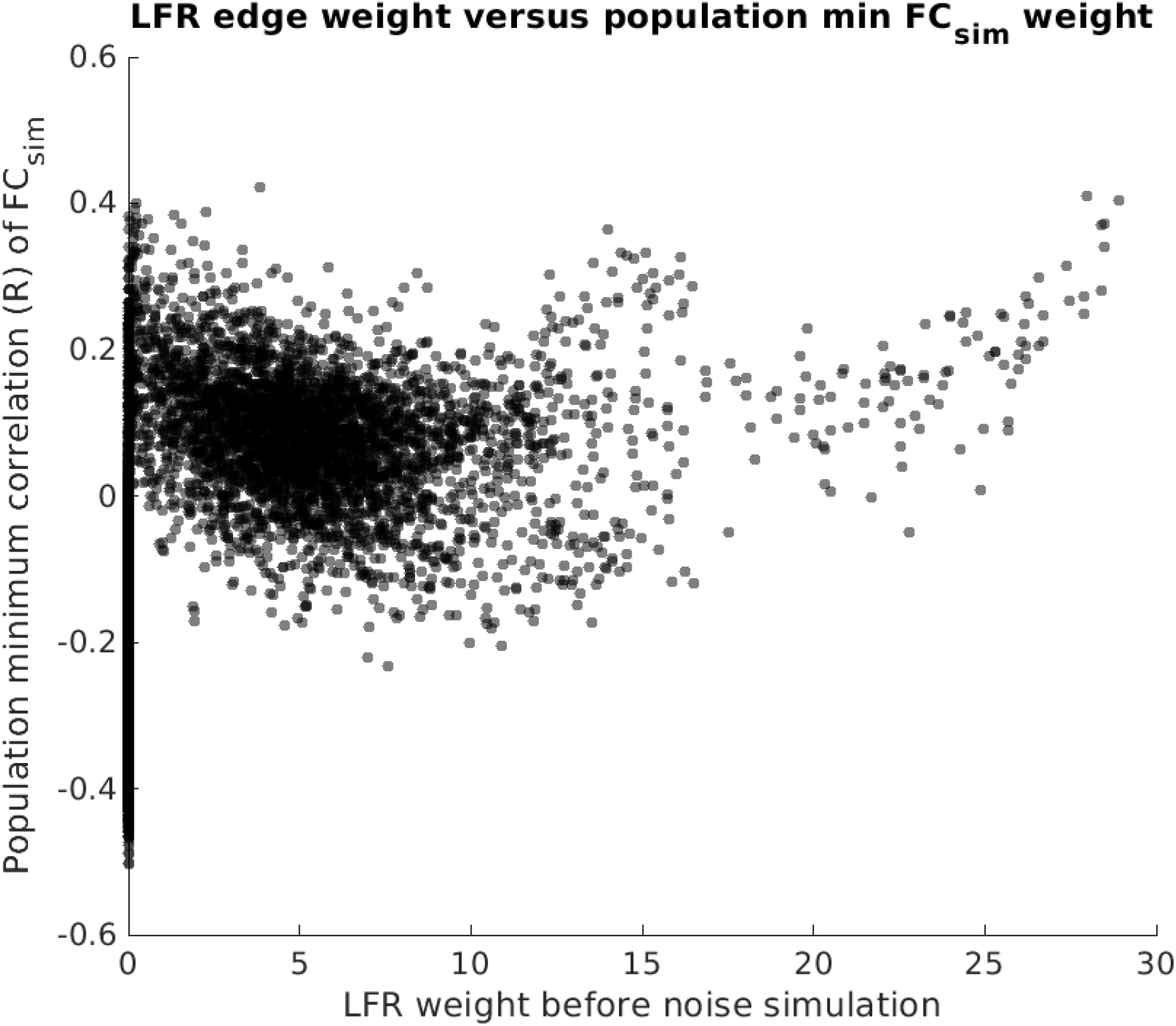
The edgewise relationship between the ground truth LFR edge weight (x-axis) and the population minimum of the corresponding FC_sim_ edge. We chose to use the population minimum, rather than the average, to depict a scenario with a high proportion of negative edges. Approximately 40% of the edges in the network are negative when considering the population minimum of each edge, and only 2.5% of negative FC_sim_ edges represent non-zero LFR edges.

Since negatively weighted edges are more likely to indicate a false positive (97.5% chance) than a true positive (2.5% chance), we reasoned that it is reasonable to zero-out the negatively weighted FC_sim_ edges. On the other hand, another common approach is to apply the absolute value function to the negative edges, thus ensuring that strongly negative edges are preserved. **Figure 4** shows that the absolute value approach will also preserve many false positives, which are the strongly negative FC_sim_ edges appearing along the y-axis. Since both removing negative edge weights and applying absolute value to them are acceptable solutions to apply to real FC data, we generated two downstream copies for each FC_sim_, one with negative weights zeroed out, and another with the absolute value function applied to the weights.

**Figure 5** depicts the histograms of the thresholds for the N=100 replicates. When considering the threshold values themselves, the objective function (using *λ*) and a GCC=1 for the M1 parameter produced a smaller threshold than the percolation method which was statistically significant (t=24.9, p<0.0001 for both absolute value; negatives zeroed out, t=224.2, p<0.0001).

**Figure 5:**
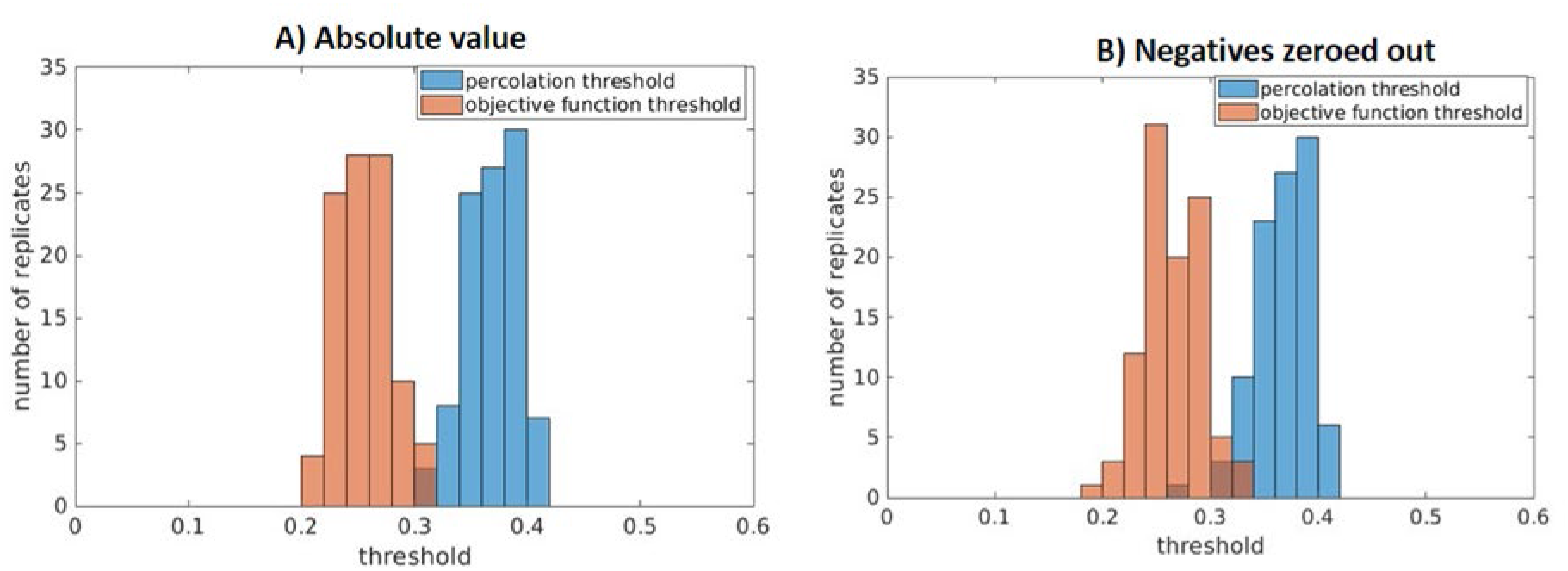
Histograms of the thresholds selected by the percolation threshold and objective thresholding methods. N=100 for each histogram, two histograms shown per panel. A. using absolute value, the average threshold for the objective function was 0.26 (standard deviation SD=0.024) and for the percolation method the average threshold was 0.37 (SD=0.023). B. when zeroing out the negative, the average threshold for the objective function was 0.26 (SD=0.028) and for the percolation method the average threshold was 0.37 (SD=0.025).

We then applied various thresholding methods such as statistical, percolation, and maximum spanning tree to each FC_sim_ network to generate a collection of binarized networks. For the OFT, we considered a variety of graph metrics as in previous work, e.g., density, *λ*, centrality measures, and found that *λ* outperformed all other choices, including density, Q, average (global) CC, efficiency, and assortativity, so we henceforth focus on results based on *λ* **(Fig 3)**. We computed the NMI between LFR community assignments, and the module assignments calculated in the FC_sim_ before and after thresholding (**Figure 6)**.

**Figure 6:.**
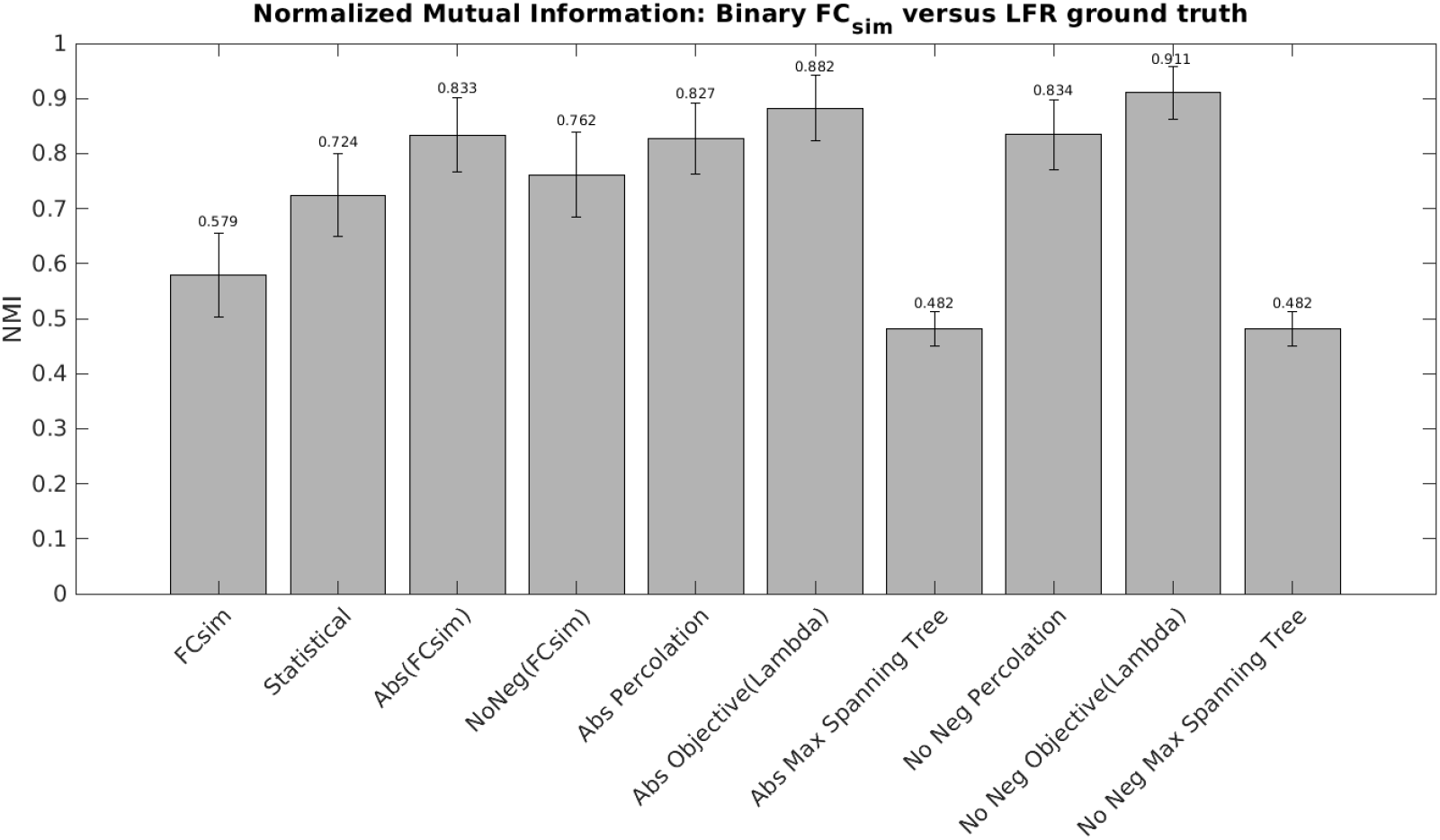
NMI comparison between thresholded FC_sim_ graphs and the LFR ground truth graph. Bar heights represent population average NMI (across N=100 replicates), and error bars represent standard deviations.

The OFT yielded the highest NMI in terms of average NMI of detected modules after thresholding to the ground truth LFR module structure, regardless of how negative weights were handled. All differences were significant, between any choice of comparisons, except for the two MST outputs (both average NMI 0.482). Although the percolation threshold and OFT applied to absolute values of correlations appear numerically close, the difference was statistically significant (t=6.46, p<0.0001). Likewise, when negative edges are set to zero, the objective function yields a higher NMI than the percolation threshold (t=11.28, p<0.0001).

Whereas NMI provides a quantitative basis for community similarity, or network organization, density provides a more straightforward perspective regarding whether the *correct number* of edges were removed (**Figure 7)**. Accuracy is defined as thresholded network density minus the density of the ground truth LFR network. A density accuracy of 0 indicates that the binarization perfectly restored network density (though not necessarily by removing the correct edges). A negative accuracy value indicates that too many edges were pruned, and a positive accuracy value indicates that not enough edges were pruned. The LFR network had a density of 0.0621. The maximum spanning tree (where density does not differ by simulation replicate) has a density of 0.0055, indicating that the LFR network has about 11 times as many edges as its maximum spanning tree. The FC_sim_ network with negatives zeroed out has a density of 0.581, indicating that 41.9% of FC_sim_ edges are, on average, negatively valued. The density of any FC_sim_ network after taking the absolute value but before thresholding would always be 1. The percolation threshold and OFT yielded density accuracy values with much smaller magnitude, with the smallest magnitudes associated with the OFT; moreover, even the closest two unequal values, derived using the percolation threshold with absolute values and OFT with absolute values are statistically significantly different (t=38.3, p<0.0001).

**Figure 7:.**
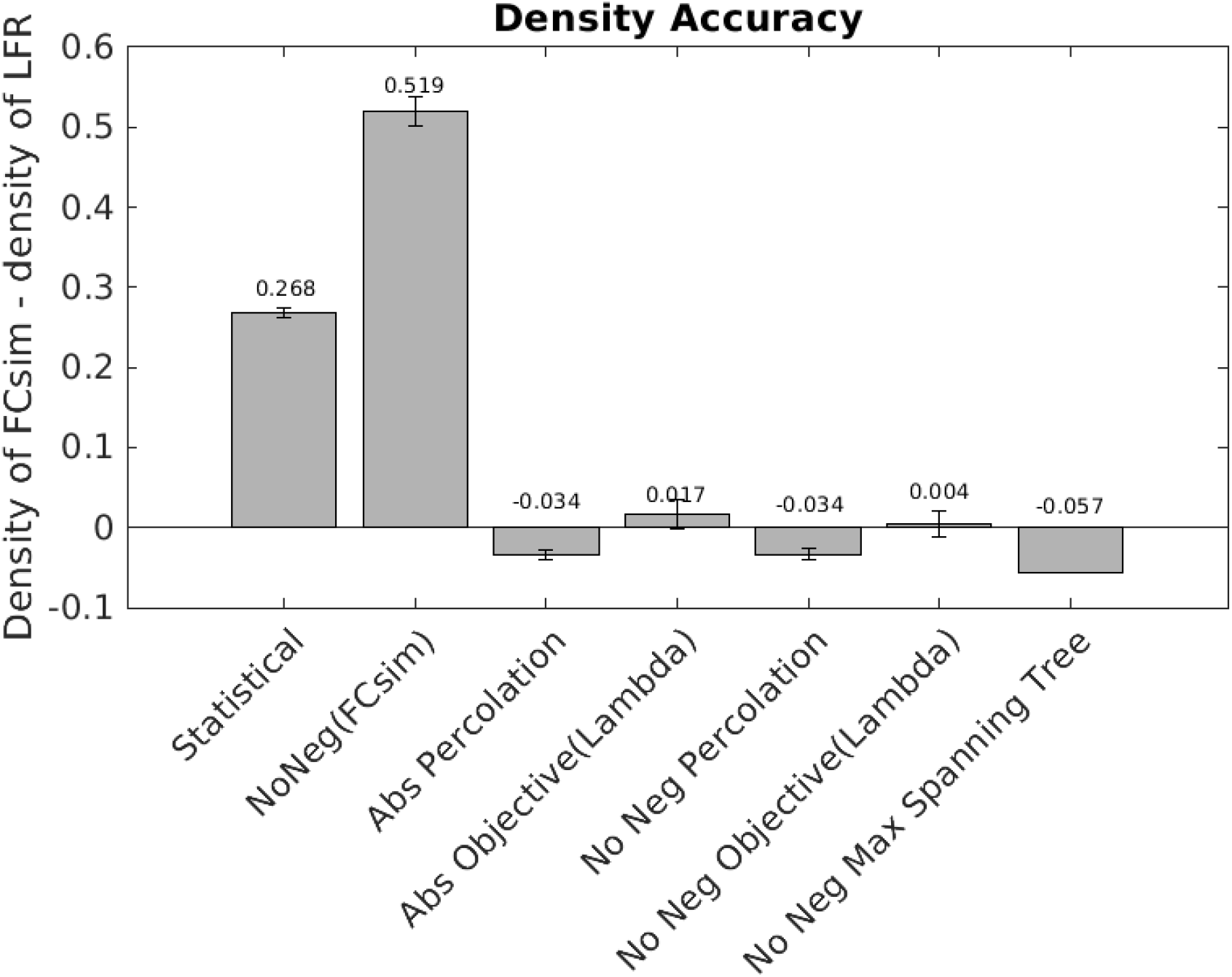
Density Accuracy. Since the LFR ground truth density is subtracted from the FC_sim_ density at each replicate, negative density accuracy values indicate that the binarization method removed too many edges, and positive accuracy values indicate that not enough edges were removed. All differences except that between the two percolation methods are significant. Bar heights represent population (N=100 replicates) averages, and error bars represent standard deviations. The maximum spanning tree has no error bars because the number of edges in the maximum spanning tree is a function of the number of nodes in the network and does not vary by replicate.

The CC is defined on a nodal basis and represents the degree to which a node’s neighbors are connected to each other. By definition, the MST has a CC=0 for all nodes. The nodal average (global) CC of the LFR network is 0.431. We defined a CC accuracy by first subtracting the LFR CC value for each node from each individual FC_sim_ replicate’s CC value for that node, then averaging the nodal accuracies across the population of replicates, and finally computing the global average over all nodes, with results illustrated in **Figure 8.** Again, the OFT significantly outperforms other thresholding methods even in the absolute value condition where the outcomes are closest (t=-10.09, p<0.0001).

**Figure 8:**
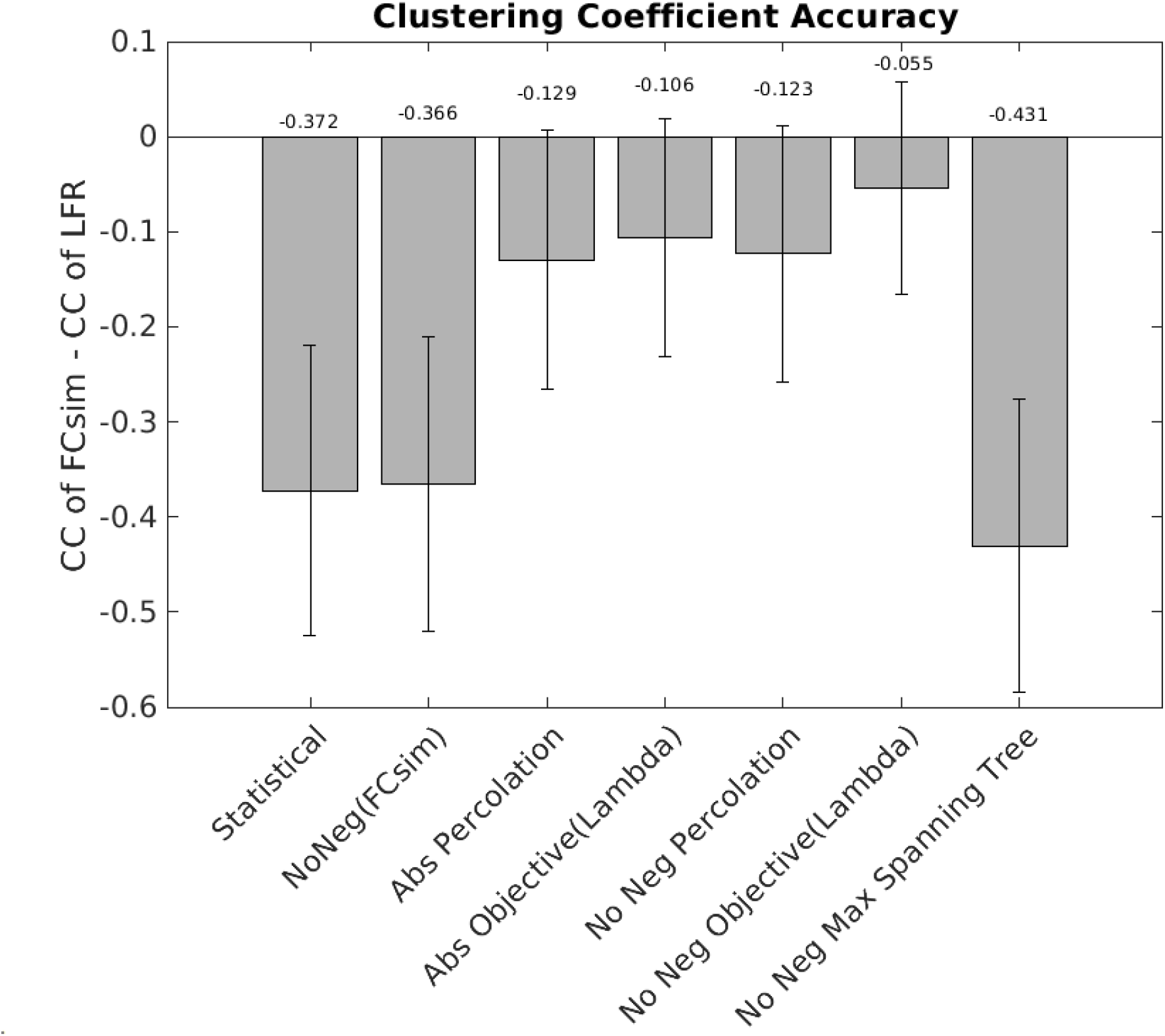
Clustering Coefficient (CC) Accuracy. The LFR ground truth CC for each node is subtracted from the CC for that node from each FC_sim_ replicate, then these nodal CC accuracies are averaged across the population of replicates, and finally a global CC accuracy is calculated by averaging over the CCs for all nodes. Negative CC accuracy values indicate that the binarization method produced nodal CC values that are, on average, lower than the nodal average CC of the LFR, and positive CC accuracy values indicate that the global CC of the binary network is higher than the global CC of the LFR. The global CC (the average of the nodal CCs) for the LFR network was 0.431. The maximum spanning tree CC accuracy has error bars because the nodes within the LFR do not have uniform CC. All differences are significant at the level of p<0.0001 after Bonferroni correction, including the differences between statistical thresholding and zeroing negatives. Bar heights represent global CC accuracy after averaging across all 100 replicates, and error bars represent standard deviations.

## DISCUSSION

Thresholding is often a critical step in network analysis with important downstream consequences. We compared several existing thresholding methods and noted drawbacks in applying these methods in some settings. To address these drawbacks, we developed an objective function thresholding (OFT) method for brain networks that can be implemented using any graph metric (**Figure 2**), and then we applied this and other thresholding methods to simulated FC networks derived from a ground truth LFR network starting point to compare the performance of these methods *in silico*. Our method is “objective” in both a mathematical sense and a semantic sense. That is, mathematical optimization is the process of finding an extremum of a function known as a cost function or objective function, which is exactly what we do to arrive at an optimal threshold value; semantically, our method is objective in that the threshold is determined by an automated procedure, without the need for subjective judgments in the process. The objective function that we use quantifies the difference between any scalar graph metric defined across a range of thresholds and its values at both endpoints of the threshold range, based on the idea that the low threshold endpoint represents an overly dense network due to the preservation of edges representing low signal and high noise, while the high threshold endpoint corresponds to an overly sparse network that lacks the redundancy characteristic of brain connectivity.

We evaluated the OFT method for a set of graph metrics (density, *λ*, efficiency, CC, Q, and assortativity) and found that *λ* was the best performing choice of metric (Figure 3). Performance was defined by the NMI between the network modules detected after binarization of simulated FC networks derived from a baseline LFR network and the ground truth modules associated with the baseline LFR network itself (Figure 1). Comparing community partitions through NMI is a widely used approach that provides a reliable measure of network community structure similarity, robust to small changes in edge-wise network content (Danon et al., 2005, Lancichinetti et al., 2009). In this work we have shown, based on NMI calculation (**Figure 6**), that the OFT method better restored the original graph structure than did previously published threshold selection methods that are evaluated, including the percolation threshold (Bordier et al., 2017), statistical thresholding (Bullmore and Bassett, 2011), MST thresholding (Kruskal 1956; Li 2022). This result held regardless of how negative values were treated in the FC_sim_ networks. Moreover, the OFT method produced node-wise CCs (**Figure 8**) and graph densities (**Figure 7**) that were closer to those of the baseline network than did the other thresholding approaches. Finally, the objective function method, by design, yielded the smallest threshold (**Figure 5**), meaning that more edges are preserved.

The OFT method is simple to apply, given that there are readily available software options for computing graph metrics. The method can be applied to any network, with any form of adjacency matrix. For example, unlike the percolation threshold, which can only be applied to path connected networks for which all nodes belong to the GCC, the objective function approach can be adapted to networks with smaller GCCs, say GCC=x, by setting the upper threshold to be the threshold at which GCC first drops to some value less than or equal to cx where c is any number in the interval (0,1] and by choosing a graph metric that is well-defined on networks that are not path connected. We find that the OFT based on *λ* outperformed the other methods across all of these performance measures and thus offers an attractive approach to consider for thresholding of brain networks based on experimental or clinical measurements. Given its intuitive underpinnings, ease of use, and superior performance in our simulation study, we conclude that the OFT method should receive serious consideration for use in thresholding in future studies of brain connectivity involving binarization of connectivity matrices.

One recent review found that “direct structural connectivity optimistically explains no more than 50%” of functional connectivity (Suarez et al., 2020), and that SC-FC edge-weight correlations range between R=0.3-0.7. In future work, OFT could be used to re-examine the relation between SC and FC data obtained from individual subjects. Such comparisons typically involve thresholding to deal with the very different types of values– non-negative integer counts of streamlines in the SC versus correlation values between - 1 and 1 in the FC– used to form networks across these two imaging modalities. As in our simulated FC data, it is challenging to handle negative values (**Figure 4**). For FC data, large negative values could be considered important, since negative correlations also represent relationships in the activity of two brain regions; hence, taking the absolute values of correlations before thresholding may be reasonable, the approach that we took in our study. Alternatively, the network can be partitioned into “negative edges only” and “positive edges only” subnetworks for separate analysis. This may create networks with GCC<1, a condition that is allowed in the OFT method. An OFT could be derived for each of these subnetworks, whereas it is likely that at least one of these subnetworks would not be path connected, which would be problematic for existing thresholding methods.

We used a quadratic objective function, which is a standard choice that avoids issues of cancellation of terms of opposite signs and provides smoothness. Other functional forms could be considered, however. In the case where *M_θ_* is monotone, the interior extremum of *F_M_(θ)* occurs at the *θ* value where *M_θ_=(M_0_+M_1_)/2*. In theory, if a practitioner had additional knowledge suggesting that the network of important edges in a dataset was in some sense more similar to M_0_ than to M_1_, or vice versa, then the weights in this average could be adjusted to yield an optimal threshold *θ* where *M_θ_*=*aM_0_+bM_1_* for some choice of *a,b* other than (1/2,1/2) with *a+b=1*.

## CONCLUSIONS

Network thresholding plays a crucial, early role in the overall process of seeking to understand the relationships encoded in brain connectivity networks. Ideally, threshold selection will be done via an automated procedure based on easily calculable quantities, to allow for reproducibility and reduce bias. The OFT method that we have proposed provides such a procedure, which can be flexibly adapted to reflect knowledge about the underlying network before it is applied, and hence represents a useful addition to our toolbox for brain network analysis.

## Abbreviations and Definitions

MRI: magnetic resonance image
ROI: region of interest
SC: structural connectome
FC: functional connectome
GCC: giant connected component
LFR: Lancichinetti-Fortunato-Radicchi
LFR-sf: the scale free LRF networks
NMI: normalized mutual information

## ACKNOWLEDGMENTS

This work was supported by grants from the US National Institute of Mental Health (NIMH) R01MH112584 and R01MH115026 (Prasad).

## AUTHOR CONTRIBUTIONS

Nicholas Theis, Jonathan Rubin and Konasale Prasad conceived the concept. Nicholas Theis conducted the analyses under the supervision of Jonathan Rubin, Satish Iyengar, Joshua Cape and Konasale Prasad. Nicholas Theis, Jonathan Rubin and Konasale Prasad drafted the manuscript. Satish Iyengar and Joshua Cape reviewed and edited the manuscript. All authors approved the manuscript.

## CONFLICT OF INTEREST

None of the authors have any conflict of interest specific to this publication.

## FUNDING STATEMENT

This work was funded through grants from the US National Institute of Mental Health (grant numbers provided above). The funding agency did not influence the contents of the manuscript.

## Notes

### Competing Interest Statement

The authors have declared no competing interest.

### Summary of Updates

The manuscript is revised with more details on the methods and the results. The discussion section has also been revised in light of the above modifications.

